# Vasomotor Symptoms Monitoring with a Commercial Activity Tracking Watch

**DOI:** 10.1101/154658

**Authors:** Darrell O. Ricke

**Affiliations:** Group 48 Bioengineering Systems and Technologies Lincoln Laboratory, Massachusetts Institute of Technology, 244 Wood Street, Lexington, MA 02421-6426

## Vasomotor Symptoms Monitoring with a Commercial Activity Tracking Watch

Continuous tracking of electrodermal activity (EDA), also known as galvanic skin response (GSR), values with commercial fitness devices for individuals with vasomotor symptoms (hot flashes) provides a path forward for future studies with fine resolution monitoring. This can improve upon the current reliance on the use of personal diaries^1^. There are multiple conditions associated with vasomotor symptoms including menopause /early menopausal transition^2^, medications (Quinestrol, tramadol, etc.), chemotherapy and Tamoxifen, hyperthyroidism, infections (Inflammatory Bowel Disease – IBD, etc.), and more. Multiple studies have characterized hot flashes in premenopausal and menopausal women using self reported methods, laboratory polysomnographic recording, and some specially designed devices^3^. However, it is difficult for an individual to track and accurately report nighttime vasomotor symptoms without the aid of a physiological monitoring device^3^. Emerging commercial and custom devices with EDA meters will greatly facilitate the nighttime monitoring of hot flashes for individuals for more informative longitudinal studies of conditions with associated vasomotor symptoms. This report illustrates the potential fine-resolution monitoring of nighttime vasomotor symptoms using commercially available activity-tracking devices with an EDA sensor.

MIT Lincoln Laboratory conducted a longitudinal study on volunteers wearing physiological monitors. The study protocol and written consent form were reviewed and approved by the MIT Committee on the Use of Humans as Experimental Subjects (COUHES). The commercial devices in the study include the Basis B1 watch and Basis Peak watch monitors for tracking heart rate, sleep with predicted sleep phases (light, deep, and REM – rapid eye movement), activity, skin temperature, and perspiration (EDA/GSR) without the use of electrodes or gel. The Basis B1 and Peak watches are no longer commercially available, but similar devices with EDA/GSR sensors are available. Vasomotor symptoms started disrupting the sleep of a woman volunteer on November 23, 2015, calling attention to their occurrence. After November 23, 2015, the volunteer started personally logging the occurrence of vasomotor symptoms, noting that the recorded EDA signals reflected the logged hot flash intensities and durations. Figure 1 shows data from eight days around this time period. Freedman^4^ identified a good agreement between an increase of 2 μS/cm in 30 s time period with patient self-reports for vasomotor symptoms. For November 15^th^, the EDA median value was 6.8e-4 μS/cm and average value of 4.9e-2 4 μS/cm. For November 16^th^, the EDA median value was 6.8e-4 μS/cm and average value of 1.9e-2 μS/cm. Across all nights tracked, the EDA median value was 3.7e-4 μS/cm illustrating baseline EDA values for this volunteer. The longitudinal data collected indicates that the volunteer’s vasomotor symptoms may have been occurring as early as June 2014, but went unnoted until higher intensity vasomotor symptoms caused sleep disruptions. A review of all sleep time data indicate an increase in sleep interruption minutes as reported by Basis (52/1957=2.9% for EDA range 10-15 μS/cm and 15/311=4.8% for EDA range 15-25 μS/cm compared to 3823/258119=1.5% for EDA < 1 μS/cm). This is consistent with volunteer observations. Nights like Nov. 23 and 24, 2015 cause sleep disruptions. Figure 2 illustrates over two years of nighttime EDA values while this volunteer was sleeping. Starting in December 2015, the volunteer started self-tracking EDA values and vasomotor symptoms. Daytime peaks were associated with both exercise and vasomotor symptoms. The volunteer reports that daytime vasomotor symptoms and sleep-disrupting vasomotor symptoms were consistent with recorded EDA peaks but they did not record these observations. Nighttime EDA peaks well above baseline values were observed in clusters from June 2014 until November 2016. Note that nights with low EDA values still occur frequently for this volunteer, indicating nights free of vasomotor symptoms. The volunteer did not take hormone or nonhormonal formulations for the treatment of vasomotor symptoms. Note that the volunteer’s Basis B1 watch was replaced with a Basis Peak watch in August 2015.

**Figure 1.**
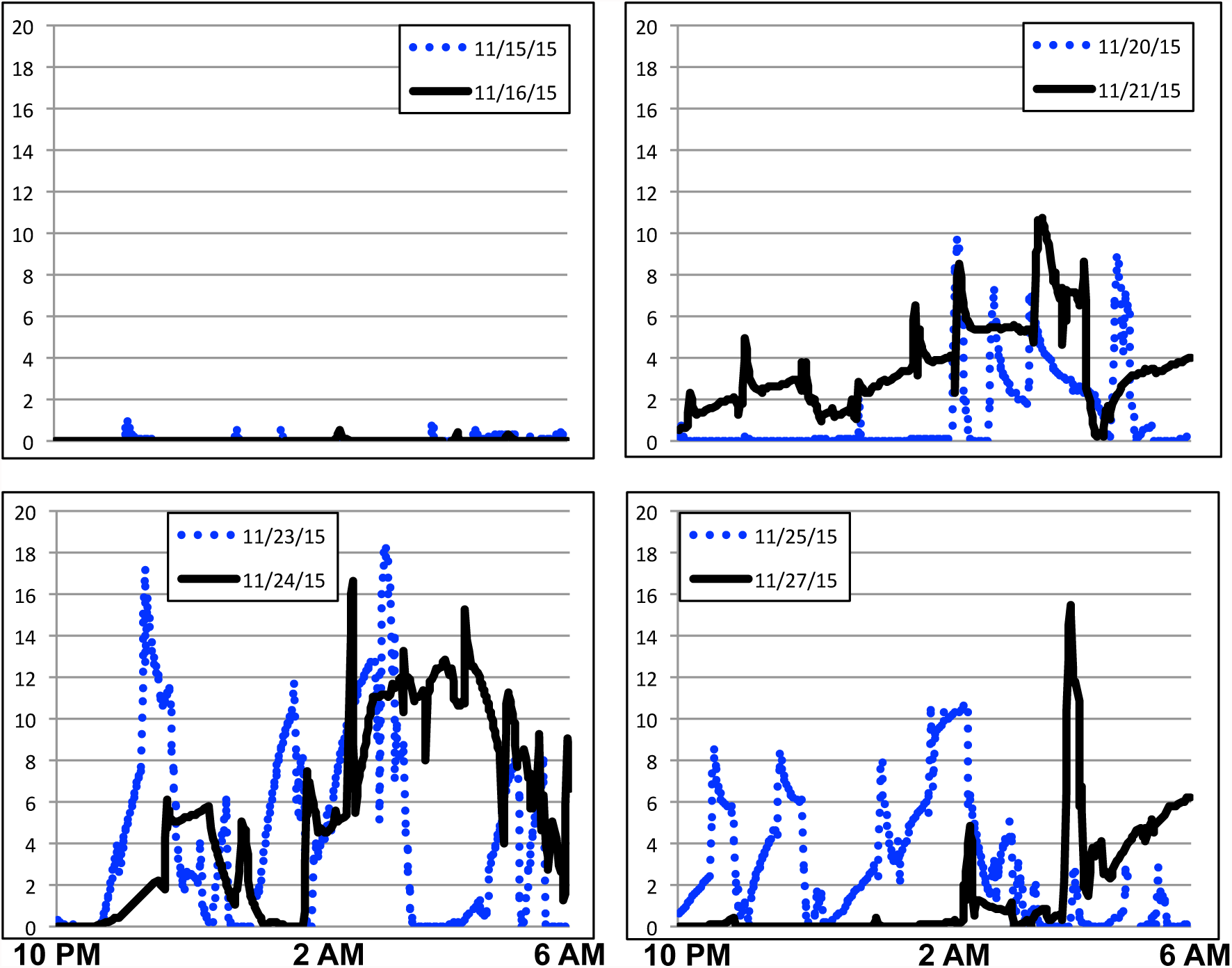
Nighttime vasomotor symptoms tracked with Basis Peak watch. Vertical axis shows EDA values in μS/cm.

**Figure 2.**
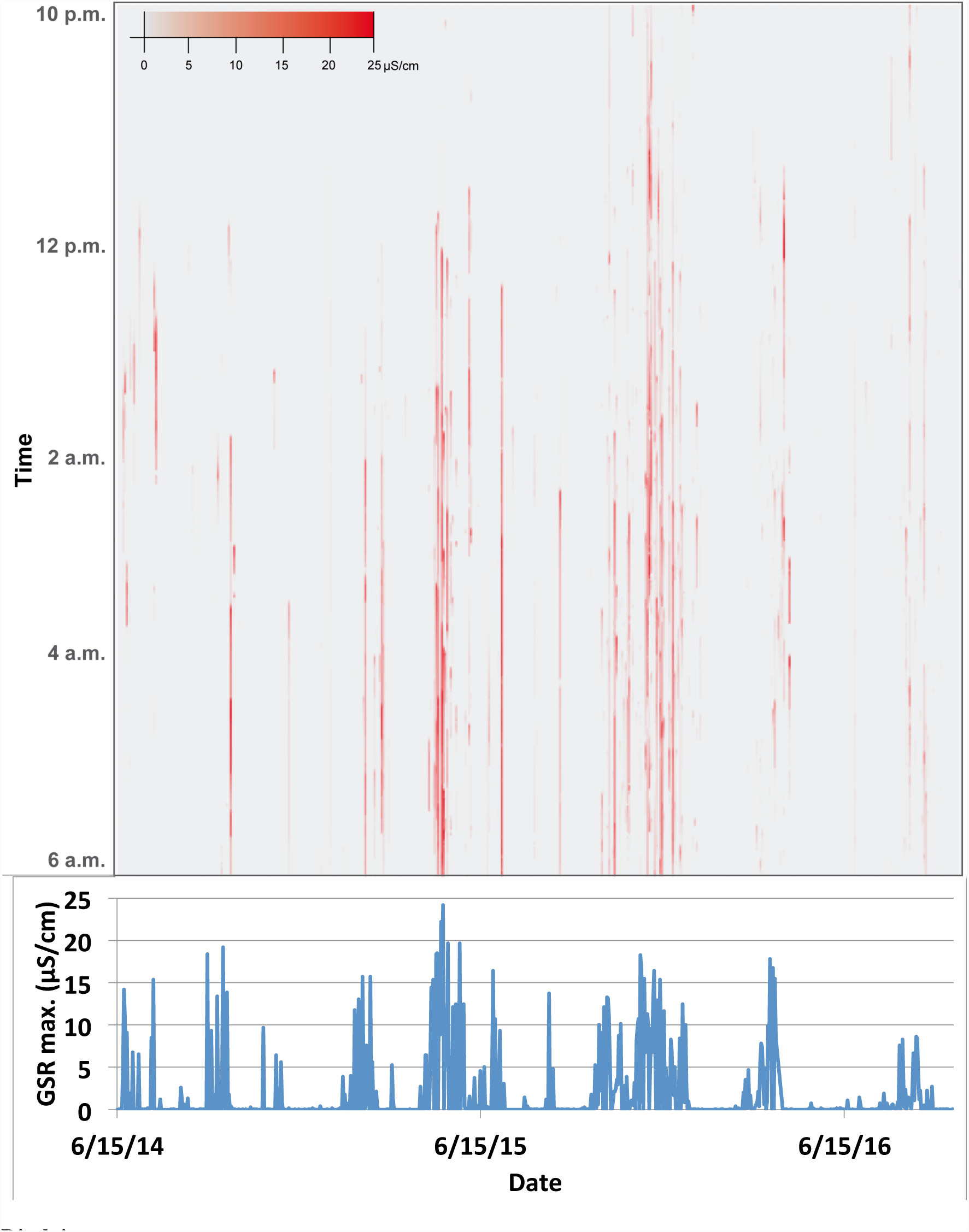
Heatmap of each minute of volunteer’s nighttime Basis EDA values tracked with Basis B1 or Basis Peak watch and line plot of nightly maximum EDA values.

Longitudinal studies of large numbers of volunteers will provide new foundations for tracking vasomotor symptoms associated with menopause and other conditions. Continuous tracking of EDA values with readily available commercial tracking devices will provide a path forward for future fine-resolution longitudinal studies of conditions associated with vasomotor symptoms. Insights into understanding and treating vasomotor symptoms will be greatly advanced by these longitudinal studies, as the different causes of vasomotor symptoms may vary in intensities and durations. EDA is also reported as a sensitive index of sympathetic nervous system activity^5^. In addition, commercial devices with EDA meters will be valuable personal monitoring tools to premenopausal and menopausal women and individuals experiencing vasomotor symptoms.

## Disclaimer

This material is based upon work supported by the Assistant Secretary of Defense for Research and Engineering under Air Force Contract No. FA8721-05-C-0002 and/or FA8702-15-D-0001. Any opinions, findings, conclusions or recommendations expressed in this material are those of the author(s) and do not necessarily reflect the views of the Assistant Secretary of Defense for Research and Engineering.

## Acknowledgements

I thank Emily Simons for graphics layout support and Paula Collins for careful review of this manuscript. Assistant Secretary of Defense for Research and Engineering has no involvement in this report.

### Contributors

DR wrote the report. Written consent to publication was obtained from the volunteer.

